# A deeper cascade of mechanisms involved in cold resistance in *Hevea brasiliensis*

**DOI:** 10.1101/2021.11.02.466997

**Authors:** Carla Cristina Da Silva, Stephanie Karenina Bajay, Alexandre Hild Aono, Felipe Roberto Francisco, Ramir Bavaresco Junior, Camila Campos Mantello, Anete Pereira de Souza, Renato Vicentini dos Santos

## Abstract

**Background:** *Hevea brasiliensis* is the main global source of natural rubber. Due to fungal disease pressure in hot, humid regions, rubber plantations have been moved to drier “escape areas” with lower temperatures. In order to analyze gene expression regulation during cold exposure, we studied young GT1 and RRIM 600 rubber tree clones with different cold tolerance strategies.

**Results:** Alongside traditional differential expression approaches, an RNA-seq gene coexpression network (GCN) was developed with 27,220 genes grouped into 205 gene clusters. The GCN related most rubber tree cold stress molecular responses to 31 clusters across three GCN modules: a downregulated group with 16 clusters and two upregulated groups with twelve and three clusters. The hub genes of the cold-responsive modules were also identified and analyzed. We observed that the general response to short-term cold exposure involves complex regulation of the jasmonic acid (JA) stress response and programmed cell death (PCD), upregulation of ethylene-responsive genes, and relaxation of florigen gene inhibition. As a result, we identified single DEGs and gained insights into the mechanisms involved in the response to cold stress in young rubber trees.

**Conclusions:** Our findings may represent the species’ genetic stress responses developed during the course of evolution, since the examined varieties were genotypes selected during the early years of rubber tree domestication. Understanding the cold response mechanisms in *H. brasiliensis* could improve breeding strategies for this crop, which has a narrow genetic base, is being impacted by climate change and is the only source for large-scale rubber production.

## 1 Background

*Hevea brasiliensis*, mainly known as the rubber tree, is an allogamous perennial tree species native to the Amazon rainforest that belongs to the Euphorbiaceae family (de Souza et al. 2016). Although rubber trees are the main global source of natural rubber (Gonçalves and Fontes 2010; Pootakham et al. 2017; De Faÿ and Jacob 2018), this tree species is a recently developed crop that is still undergoing domestication (Priyadarshan and Clément-Demange 2004). The dispersal and domestication of rubber trees around the world was based on only approximately 20 seedlings that were introduced in Southeast Asia in the late 19th century. These seedlings were the only survivors of a collection of thousands of seeds from the Amazon Basin, resulting in the selection of elite trees and controlled hybridization for reproduction in Hevea. Thus, to date, almost all commercial varieties of *H. brasiliensis* planted are derived from these seedlings; therefore, their genetic variability is quite narrow (Gonçalves and Fontes 2010; de Souza et al. 2015).

Although the Amazon Basin offers ideal conditions for rubber tree cultivation, the occurrence of fungal South American leaf blight (SALB) disease in the region hinders the establishment of rubber tree plantations in the area. To overcome this situation, Brazilian rubber tree plantations were moved to suboptimum or escape areas such as the Brazilian Center and Southeast regions, which are drier and present lower temperatures during the winter (Priyadarshan et al. 2009). Southeast Asian countries, which have similar Amazonian climate conditions and successfully avoided the introduction of SALB, are major rubber producers worldwide. Nevertheless, these Asian plantations have already suffered heavy losses due to other fungal disease outbreaks (Priyadarshan and Goncalves 2003; International Rubber Consortium Limited (IRCo) 2019; Pornsuriya et al. 2020) and are being relocated to suboptimal areas as well, such as Southern China and Northeast India (Priyadarshan et al. 2009). Thus, rubber tree varieties with adaptations to areas with new edaphoclimatic conditions are of primary interest.

Among the stress factors characteristic of the escape areas, low temperatures heavily affect *H. brasiliensis* development, causing damage to leaves and latex production (Priyadarshan et al. 2005). For a rubber tree plantation, a prolonged period of low temperatures or a frost occurrence may cause plant death (Priyadarshan et al. 2009). Cold stress triggers massive reprogramming of plant gene expression to adjust metabolic processes to cope with the cold environment (Theocharis et al. 2012). To promote breeding to select for cold tolerance in *H. brasiliensis*, a deeper understanding of the species cold response mechanisms is necessary.

In recent years, due to the impact of low temperatures on rubber tree plantations and rubber production, there has been an increase in the number of studies on *H. brasiliensis* molecular cold stress responses, especially using comparative transcriptomics (Silva et al, 2014; Deng et al. 2018; Gong et al. 2018; Mantello et al. 2019). Examining the modification of Hevea gene expression under cold treatments has already enabled the identification of thousands of differentially expressed genes (DEGs) (Cheng et al. 2018; Deng et al. 2018; Sathik et al. 2018). However, although the comparative transcriptome analysis of resistant and cold-sensitive clones has shown that cold is a key factor in latex production, more precise information on the main biological pathways and metabolic processes involved in the cold response of rubber trees has not yet been obtained. Recently, Ding et al. (2020) modeled a gene coexpression network (GCN) and associated a network module with cold treatment. Although the cold response mechanisms were not assessed, the GCN created represented a rich source of data for investigating cold response.

Here, we carried out a study based on a GCN modeled with transcriptomic data from two of the earliest *H. brasiliensis* clones to identify novel patterns in the genes involved in the response of *H. brasiliensis* to cold stress, aiming to gain deeper insight into how young rubber tree plants cope with low temperatures. The use of the transcriptome from a cold experiment with two genotypes, one resistant and one susceptible, allowed the identification of the central genes in the created networks underlying cold stress in rubber trees. Thus, this study is one of the pioneers in assessing the association of molecular mechanisms of Hevea through GCNs, being the first to unveil rubber tree cold stress responses in important commercial varieties.

## 2 Results

### 2.1 Bioinformatics and Differential Expression Analysis

Approximately 530 million paired-end (PE) reads were obtained for the RRIM600 genotype and 633 million for the GT1 genotype. After quality filtering, approximately 933 million PE reads were retained and used to assemble the transcriptome in Trinity software. A total of 104,738 isoforms (51,697 genes) were identified with sizes between 500 bp and 22,333 bp, with an average of 1,874 bp and N50 of 2,369. From these genes, we identified a total of 74,398 unique proteins related to 9,417 different GO terms (Table S1). The DEG analyses resulted in a final number of 30,407 genes that, analyzed through PCA, revealed a distinct profile of samples belonging to each different genotype (Fig. 1a - Principal component analysis (PCA) scatter plot for the first two principal components (PCs) of the rubber tree transcriptome data (GT1 and RRIM600 genotypes). Each point represents a different sample, colored according to the respective genotype).

**Fig. 1.**
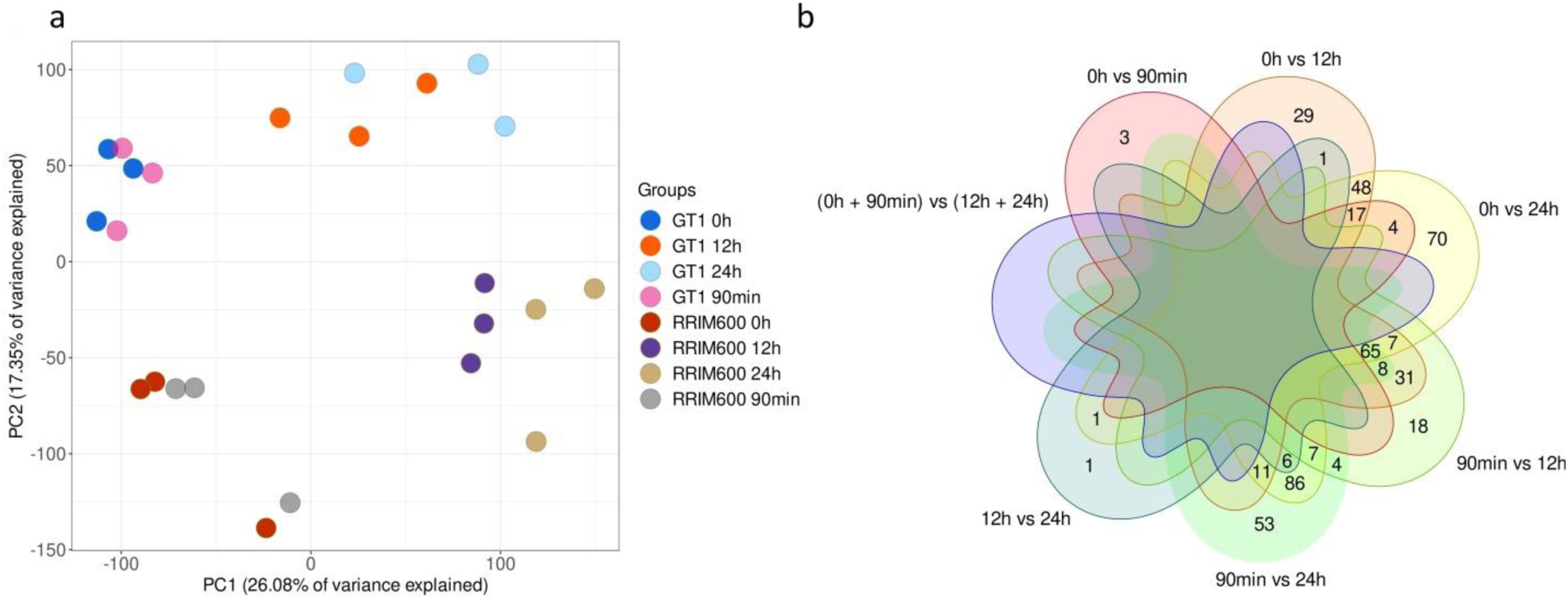
(a) PCA plot for the first two principal components of the rubber tree transcriptome data

The number of DEGs was different in each period comparison (Table 1), as there were different profiles of genes associated with each period analyzed (Table S2). Although there were apparent intersections among the tests established (Fig. 1b - Venn diagrams of differentially expressed genes (DEGs) identified under the established conditions), a joint mechanism of several cold response genes was clearly observed. We were able to annotate approximately 93% of the downregulated genes and 96% of the upregulated genes. Regarding the GO enrichment analysis, the full DEG set was enriched for response to stress and response to chitin terms. The downregulated group of genes did not present any overrepresented category. In contrast, the upregulated gene group was enriched for the following terms: response to chitin, defense response, response to wounding, cell death, photoprotection, calcium ion transmembrane transporter activity and integral to plasma membrane.

**Table 1.** Number of *Hevea brasiliensis* differentially expressed genes (DEGs) that were modulated by cold stress.

| Time period comparison | DEGs | Upregulated DEGs | Downregulated DEGs |
| --- | --- | --- | --- |
| 0 h vs. 90 m | 32 | 32 | 0 |
| 0 h vs. 12 h | 673 | 558 | 115 |
| 0 h vs. 24 h | 1,208 | 852 | 356 |
| 90 m vs. 12 h | 526 | 455 | 71 |
| 90 m vs. 24 h | 917 | 700 | 217 |
| 12 h vs. 24 h | 18 | 15 | 3 |
| (0 h + 90 min) vs. (12 h + 24 h) | 2,436 | 1,575 | 861 |

After 90 minutes of cold treatment, only thirty-two transcripts were characterized as DEGs, all of which were upregulated during the plantlets’ early response to cold stress. Of these, 26 genes were successfully annotated (Table S2). The annotated DEGs are known to be induced by cold as well as other abiotic stressors, such as drought and salt stresses. They are involved in cell wall tightening processes, growth inhibition, oxidative stress protection, calcium (Ca^2+^) signaling and cold-induced RNA processing. The late response, considered the interval between twelve hours and 24 hours of cold treatment, contained 18 DEGs, including genes that stimulate an increase in the membrane diffusion barriers and cell wall modifications.

One hundred thirty-eight DEGs were annotated with the GO term DNA-binding transcription factor activity, with 103 upregulated and 35 downregulated. All but one of them were grouped in the cold stress response gene clusters (see below). An analysis of overrepresented GO terms in the 138 DEGs showed that they are involved in regulating cellular metabolic processes: negative and positive regulations are modulated by the up and downregulated groups, respectively. While the upregulated genes are responsive to stress and other stimuli, the downregulated genes promote plant development and growth. All transcripts annotated as members of the Apetala2/Ethylene-responsive transcription factor (AP2/ERF) family and that were identified as DEGs (38) were upregulated. DEGs annotated as transcriptional repressors of jasmonic acid (JA)-mediated responses (i.e., TIFY/JAZ proteins) were also upregulated.

Nonspecific serine/threonine protein kinases (EC: 2.7.11.1) were the most highly represented enzymes among the annotated DEGs: 165 were upregulated, while 57 were downregulated. The upregulated kinases had an overrepresentation of GO terms related to reproduction processes, response to abscisic acid (ABA) stimulus, calmodulin binding and signal transmission, while the downregulated kinases were enriched for posttranslational protein modification. The second most differentially expressed enzyme types were RING-type E3 ubiquitin transferases (EC: 2.3.2.27), with 42 and 11 genes up- and downregulated, respectively. RING E3 ubiquitin proteins are essential for the ubiquitin proteasome system (UPS) due to their roles in different developmental processes and stress responses, as these proteins recruit substrates to be degraded (Cho et al. 2017). In addition, enzymes that attenuate the JA-mediated stress response were upregulated. Interestingly, enzymes that synthesize JA precursor molecules (Oxylipins) were upregulated.

### 2.2 Rubber Tree GCN Analysis

From the transcriptome data for the GT1 and RRIM600 genotypes, an HRR network was modeled, composed of 27,220 nodes across 50,650 connections (edges), to represent the cold response of the species (Fig. 2a – Modelled gene coexpression network (CGN); each node represents one gene, colored according to the differential expression analysis). The major connected component within the network is composed of 25,608 nodes (94.1%). There were also 626 other components for the remaining 1,592 genes, with a gene number within each component ranging from 2 to 11 nodes. To identify putative gene associations within the network structure, we analyzed the network with a heuristic cluster chiseling algorithm (HCCA), identifying 832 clusters (Fig. 2b – GCN); each node represents one cluster, colored according to the presence of differentially expressed genes). From these sets of genes, 626 corresponded to the disconnected components (mean cluster size of ∼3), and 206 corresponded to the core GCN structure (mean cluster size of ∼124), comprising clusters with sizes ranging from 45 to 280 genes.

**Fig. 2.**
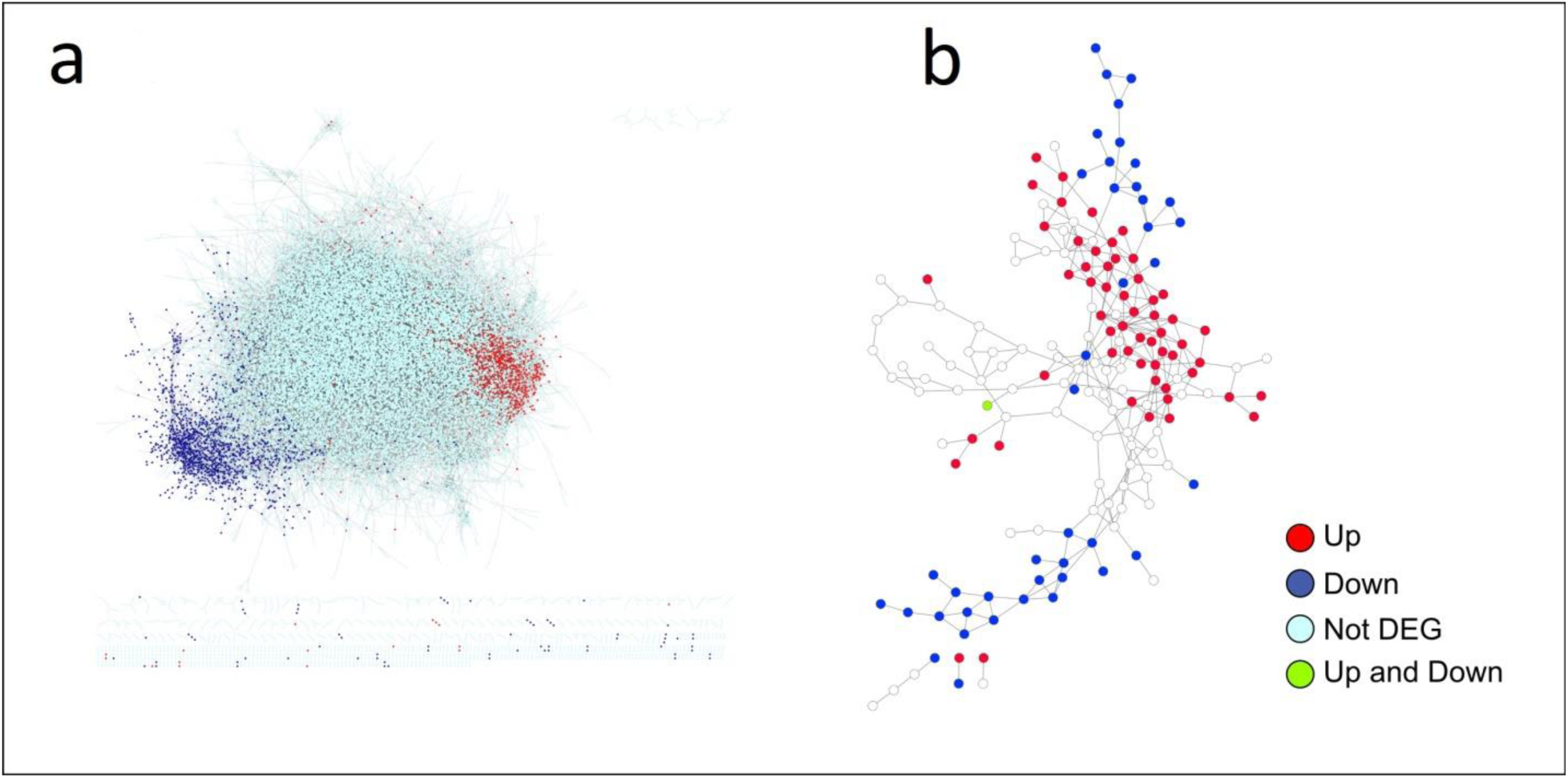
(a) CGN modeled and (b) GCN colored according to the differential expression analysis

From the 206 main clusters, 39 (14%) had GO terms that were significantly enriched, as highlighted in Fig. 3a (Fig. 3a – Modeled gene coexpression network (CGN); each node represents one gene, colored according to the enriched Gene Ontology (GO) term identified in the related cluster) for biological process GO terms. Response to stress was the most overrepresented term among clusters, encompassing 1,039 genes across 9 groups, which was expected considering the cold stress experimental design. Two of these clusters (c14 and c28) were enriched for developmental processes as well, indicating the tight correlation between environmental stimulus and growth. Interestingly, we also observed seven clusters enriched for virus replication and transposition.

**Fig. 3.**
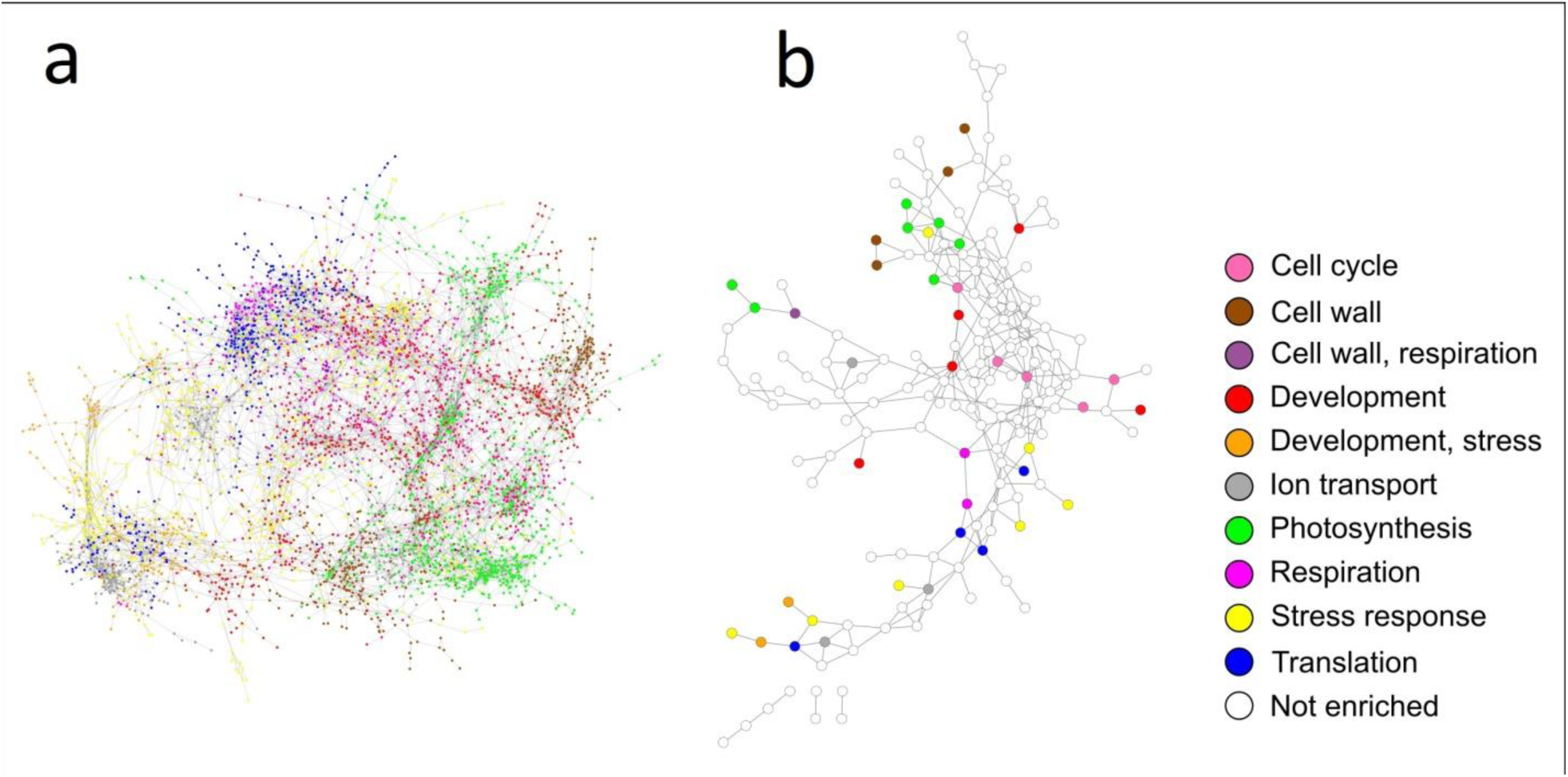
(a) CGN modeled and (b) GCN colored according to the enriched GO terms

The gene centrality measures (Table S3) showed that of the total of 27,220 genes inserted into the network, there were only 507 genes with the maximum permitted number of connections (ten). One of them, annotated as “probable cytochrome c biosynthesis protein” (CCBS), also presented the highest values of SC and BC centralities, indicating that this gene exerted great influence in the network. The gene belonged to cluster c8, which was enriched for photosynthesis but also for cellular respiration and transposition. By evaluating the intercluster connections in the HCCA-based contracted network (Fig. 3b - GCN according to the groups identified by the HCCA); each node represents one cluster, colored according to the enriched GO term; and Table S4), we found that the cluster with the greatest number of connections was c117 (nine edges), which was enriched for developmental processes.

### 2.3 Gene Clusters Mostly Related to Cold Stress Response

Even though the DEGs identified were dispersed among the GCN (Fig. 2a), specific patterns could be observed in several clusters. It was possible to identify 1,751 DEGs involved in clusters associated with the cold stress response (Table S5) (Fig. 4a Gene coexpression network (GCN) colored according to the modules considered to be associated with the cold response.). Expanding the module across these clusters’ neighbors, we established three GCN modules for the 31 clusters selected: (i) a downregulated group with 12 clusters, (ii) an upregulated group with 11 clusters, and (iii) an upregulated group with 3 clusters (Fig. 4b GCN according to the groups identified by the heuristic cluster chiseling algorithm (HCCA); each node represents one cluster, colored according to modules selected and sized according to the proportion of DEGs.). Even though there were additional clusters containing DEGs, only the most pronounced ones were used for novel inferences. Among the DEGs in the selected clusters, 480 genes were annotated with enzyme code (EC) numbers, with 339 upregulated and 141 downregulated transcripts, and 180 DEGs were assigned to KEGG pathway maps.

**Fig. 4.**
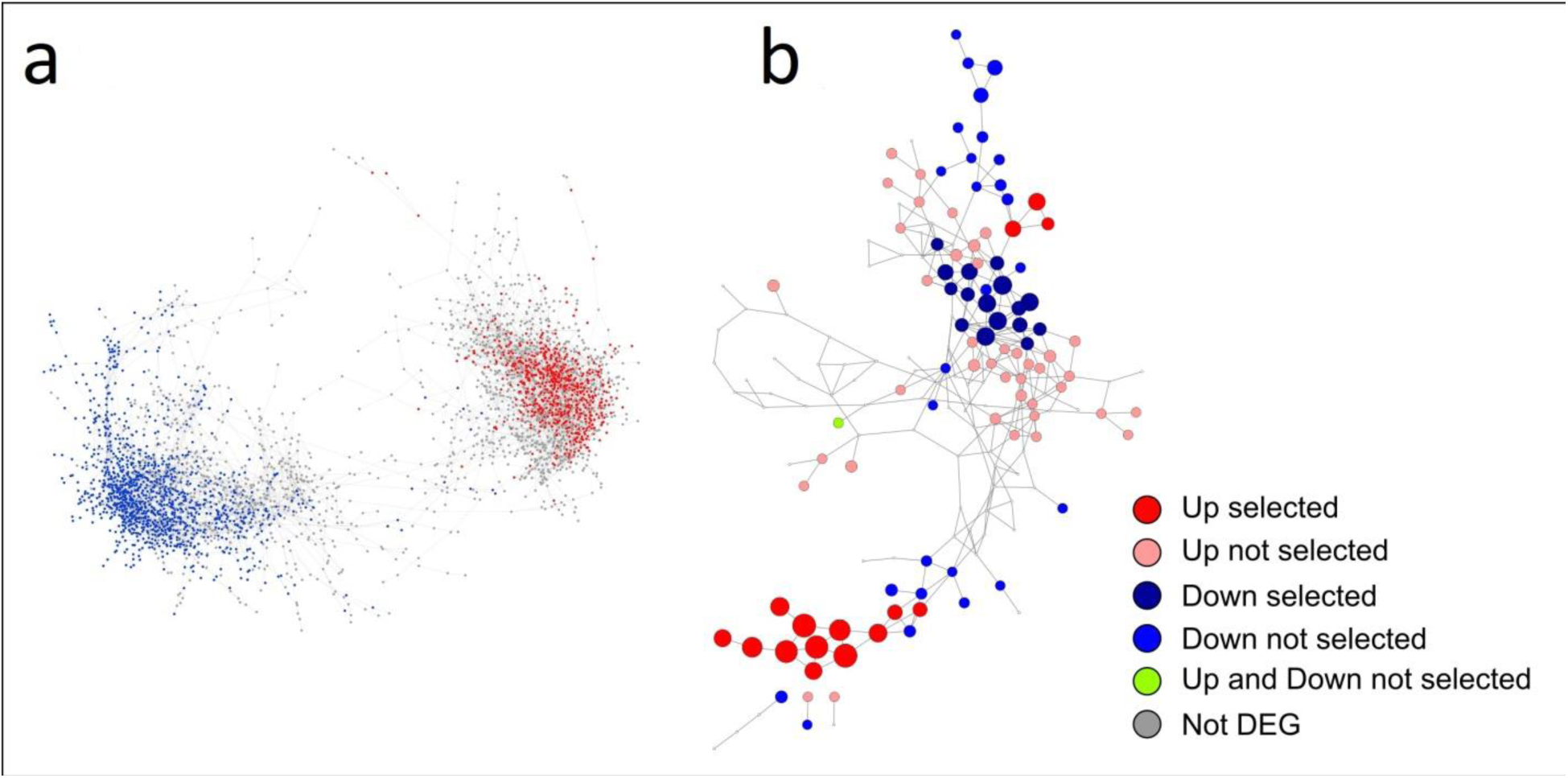
(a) GCN showing cold response modules (b) GCN showing modules selected/sized by the proportion of the DEGs

The downregulated module (i), as a whole, did not present any overrepresented GO category; however cluster c68, by itself, was enriched for the cell cycle. In this module, we identified nineteen hub genes. Almost all hubs were late-responsive genes. Among the hubs, there were genes similar to proteins that inhibit the flowering process and the ABA signaling pathway (Table S6).

The GO enrichment of module (ii) showed that signaling processes were amplified since MAP kinase kinase kinase and calcium ion (Ca^2+^) transmembrane transporter activities were overrepresented. The group was also enriched for response to stress, photoprotection and plant-type hypersensitive response. Additionally, we also observed in (ii) clusters enriched for specific GO terms. Twenty-five early-response genes were grouped into cluster c14, which was the cluster with more overrepresented GO categories among the groups of the modules selected, enriched for response to high light intensity, response to reactive oxygen species and transcription regulation activity. This group harbors several *ERF* genes, some of which are early-responsive to cold stress. Only six enzymes were early responsive to the cold treatment, and all were placed in this cluster. Four of them were annotated as RING (really interesting new gene) -type E3 ubiquitin transferases. Other clusters from this module were enriched for stress response (c28, c126 and c159), ion transport (c34), development (c28) and translation (c143).

Cluster c159 included transcripts annotated as ERFs, calmodulin binding proteins (CBPs), serine/threonine protein kinases and disease-responsive genes. Three early-responsive transcripts were present in this cluster as well. Clusters c53 and c143 included transcripts annotated as proteins involved in protein phosphorylation, serine/threonine kinase activity, stress-responsive TFs, nucleotide sugar biosynthesis enzymes and cell wall biogenesis proteins. Among the 108 transcripts that had a functional annotation in cluster c28, 25 (23%) were TFs involved in the regulation of plant defense response and development. For cluster c126, there was also an overrepresentation of GO terms for phenylalanine biosynthesis and sphingolipid biosynthesis processes. Several genes annotated as CBPs and Ca^2+^ transporters were grouped into cluster c34. Although clusters c101, c121, c161 and c194 were not enriched for any GO term, these clusters included genes involved in the ethylene biosynthesis pathway, plant development, stress response, PCD and lignin biosynthesis.

Eighteen hubs were identified in module (ii), three of which were early-responsive genes. Some hubs were annotated as proteins involved in lignin biosynthesis and stress response, while four were annotated as uncharacterized proteins. Interesting, the second transcript with the highest BC and SC values in the whole network was in cluster c28 and did not have a functional annotation. It had seven connections, four of which had genes from different clusters. Its sequence was conserved among plant species, and it was described as an uncharacterized protein.

The genes clustered in the second upregulated module (iii), according to the enrichment analysis, were mostly involved in carbohydrate metabolic processes, lipid phosphorylation and phosphatidylinositol phosphate biosynthetic processes. By itself, cluster c106 was enriched for developmental growth and water homeostasis. Three late-responsive and two non-DEG genes were identified as hubs in module (iii).

The GO terms identified in the modules selected were classified into several biological processes, separated into upregulated (Fig. 5a) and downregulated (Fig. 5b) associated categories. Although upregulated and downregulated DEGs from these modules presented evident similarities in the GO visualization, we clearly observed that the same biological processes were affected in specific ways in the upregulated modules (Fig. 5a) and downregulated modules (Fig. 5b).

**Fig. 5.**
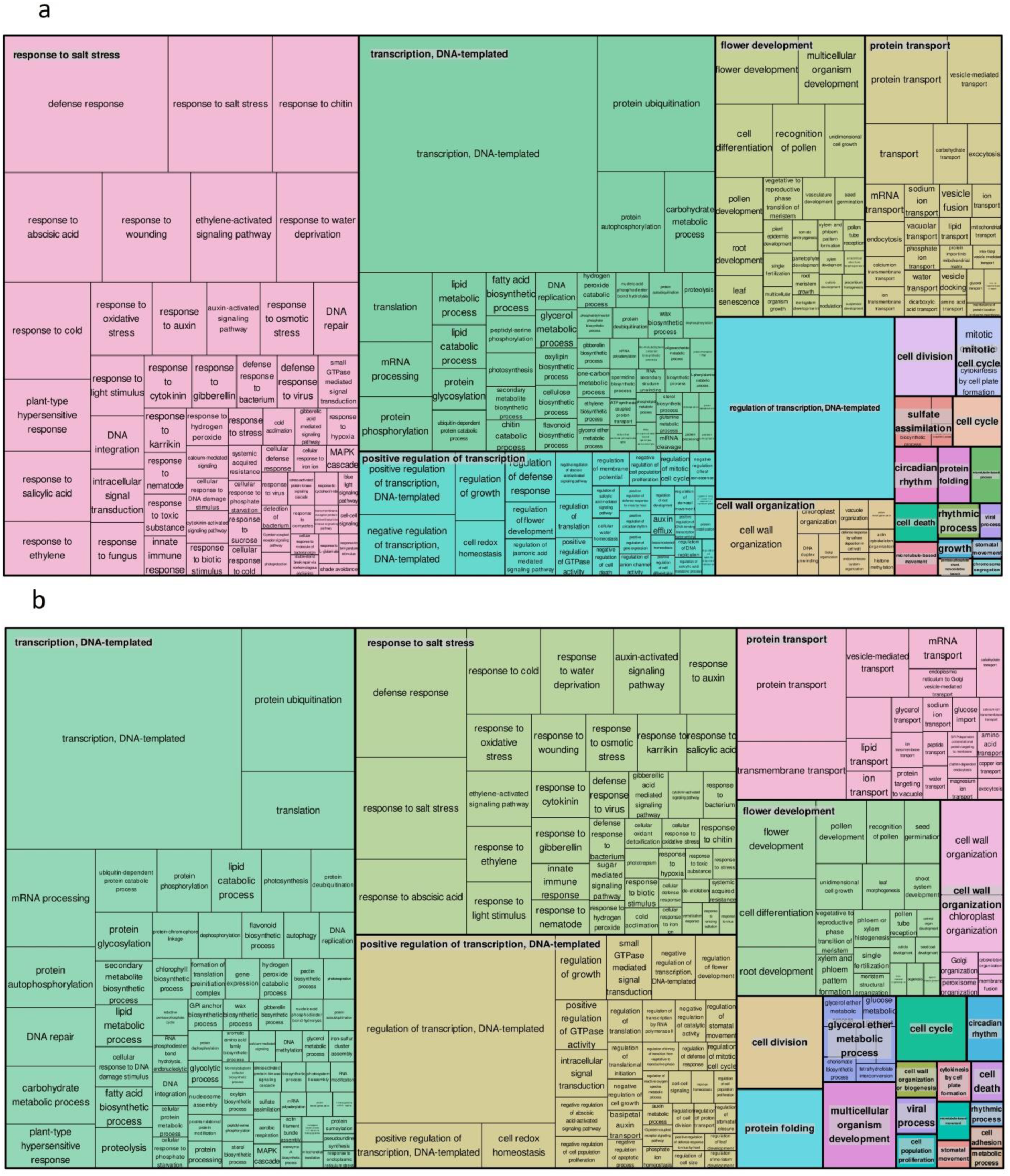
Gene Ontology terms for the cold response modules selected: (a) Upregulated modules; (b) Downregulated module

Within the upregulated groups, cold exposure also triggered the upregulation of enzymes involved in the synthesis of protective photosynthetic pigments, such as zeaxanthin and lutein (c106) and lipoic acid (lipoamide) (c194). Several enzymes from the nucleotide sugar metabolic pathway were upregulated (c101, c106, c121, c130, c143, c16, c161, c34, c53), as well as enzymes from the methionine metabolism pathway that produce ethylene as their final product (c121, c28, c34). α-Linolenic acid metabolism enzymes that synthesize oxylipins were members of clusters in module (ii).

## 3 Discussion

The optimal conditions for *H. brasiliensis* development are temperatures between 22 and 30 °C, relative humidity of 70% or higher, and annual rainfall between 1,500 and 3,000 mm. Currently, rubber tree plantations are mostly located in Southeast Asia, which has ideal edaphoclimatic conditions for crop development. Interestingly, such varieties are genetically closer to genotypes derived from the southern part of the Amazon basin (de Souza et al. 2015), also exhibiting an adaptation capacity to Amazon rainforest dynamics. However, in Brazil, the premiere region of natural rubber productivity, the state of São Paulo, is found in suboptimal conditions. Cold weather and fluctuations in rainfall levels directly affect plant development, impacting rubber production (Rao and Kole 2016). In this context, Brazilian rubber tree breeding programs have focused on the development of genotypes with high productivity and tolerance to the stress imposed by these areas of higher productivity, especially cold, which inhibits biochemical reactions, reduces photosynthetic capacity and alters the permeability of the plant membrane (Sage et al. 2008; Sevillano et al. 2009; Mai et al. 2010).

The adjustment and adaptation of the rubber tree to the climate of its current places of production depends on its acclimatization capacity. This process is essential for the development of plants that exhibit tolerance to freezing when exposed to low temperatures (Thomashow 1999). The expression of specific genes is altered at the transcriptional level during the cold acclimation process, forming several cooling-responsive pathways from a signaling network (Thomashow 1999; Carvallo et al. 2011; Jeon and Kim 2013), which we were able to clearly visualize in the combined GCN of two of the earliest rubber tree clones, GT1 and RRIM600; RRIM600 is the most planted clone in Brazil. GT1 is a primary clone obtained through open pollination of wild seedlings introduced in Southeast Asia and their unselected progeny. RRIM600 was selected in the first rubber tree breeding cycle, when breeders first used controlled pollinations. Both genotypes are prime progenitors of modern rubber tree clones (Priyadarshan et al. 2009). Considering the long life cycle of rubber tree and the few generations that occurred, these clones are substantially close to their wild ancestors; therefore, they may be considered genotypes bearing natural genetic variation in wild Hevea. Thus, the work carried out in this paper, which started from transcripts of genotypes that represented a good part of the genomic characterization of rubber trees, promoted our understanding of the response to cold in the species and which biological pathways to focus on the next steps of rubber tree breeding.

The understanding of the global cold response mechanism of the species is important for advancing the improvement of rubber production. Our results showed that there were differentially expressed genes that act in a convergent strategy to cold stimulation, considering all transcriptional triggering of GT1 and RRIM600 when exposed to cold.

The identification of genes modulated in response to cold showed that there were contrasting amounts of DEGs according to the time of exposure to cold. The comparison between 0 h and 24 h stages was the most informative in terms of number of DEGs, while 0 h and 90 min was the interval that had the lowest number of DEGs, suggesting that the seedlings were able to withstand a short period of cold exposure without massive activation or repression of genes. After twelve hours of exposure to cold, we observed that both genotypes underwent reprogramming in their expression profiles (DEGs identified at 0 h vs. 12 h), and the new changes were nearly completely maintained over the next twelve hours (DEGs identified at 12 h vs. 24 h).

The most prominent transcription factors among the DEGs were members of the *ERF* family. *ERFs* are activated by the ethylene signaling pathway and regulate the transcription of ethylene-responsive genes, which negatively impact plant growth (Müller and Munné-Bosch 2015; Van den Broeck et al. 2017). The *ERF* family in rubber tree comprises 115 members (Duan et al. 2013), and the overexpression of an Arabidopsis *ERF1* ortholog in rubber tree plants enhanced their tolerance to abiotic stresses and augmented laticifer cell differentiation (Lestari et al. 2018). In our analysis, all DEGs annotated as ERFs were upregulated, suggesting that *H. brasiliensis* plantlets maintained the ethylene-mediated stress response during the entire experiment. Additionally, ethylene biosynthesis enzymes were upregulated, indicating that ethylene plays a constitutive role in the rubber tree cold stress response.

JA biosynthesis and signaling pathways appeared to be toned down in the seedlings during the 24 h of cold treatment. *TIFY*/*JAZ* TFs, which were upregulated in our experiment, are specific to plants and are negative regulators of JA-induced responses. Nevertheless, their expression is induced by JA, which creates a regulatory feedback loop for fine-tuning of JA signaling (Chini et al. 2007). Different types of stresses induce the expression of these genes, and they are essential for plant growth-defense balance (Zhu 2016; Major et al. 2017; Guo et al. 2018; Xu and Weng 2020) Accordingly, enzymes that attenuate JA-mediated responses, such as IAA-alanine resistant 3 (IAR3) and cytochrome P450 94C1 (CYP94C1) [that catalyze the turnover of JA (Heitz et al. 2012; Widemann et al. 2013), were upregulated as well. Despite the attenuation of JA responses, the biosynthesis of oxylipins appeared to be promoted since the expression of their biosynthetic enzymes was upregulated, suggesting that rubber trees employ oxylipins rather than JA to trigger responses to short periods of cold stress. Oxylipins have their own signaling pathways, being regulators of several JA-responsive genes (Taki et al. 2005). These molecules are involved in the regulation of plant development, promote ABA biosynthesis (Dave et al. 2016), act cooperatively with ABA to control stomatal aperture during drought stress (Savchenko et al. 2014) and are promoters of induced systemic resistance in maize (Wang et al. 2020).

In addition to the findings with DEG analyses, we expanded the inferences on cold response using GCNs. Considering that cold tolerance is the result of a network of complex biochemical pathways, an appropriate way to understand such regulation is not to evaluate a single gene but a set of connected genes corresponding to a broad molecular mechanism. The correlation of how the coordinated expression of genes involved in multiple metabolic pathways act in response regulation can be elucidated through the definition of clusters within the GCN (Umer et al. 2020; Sun et al. 2021). In this way, GCN is a promising approach to better elucidate the participation of genes linked to cooling stress.

Recently, Ding et al. (2020) modeled a GCN using several public rubber tree transcriptomes, providing insights into rubber biosynthesis. Despite their great potential, such approaches are incipient in rubber tree research, and this study is the first to evaluate the *Hevea* cold response from a more holistic perspective.

### 3.1 Selection of Cold Response-Associated Clusters

The selection of GCN clusters associated with the cold response allowed us to supply a range of molecular mechanisms triggered by such a stress. The starting point of the rubber tree cold stress response mainly resided in cluster c14. Several *ERF* genes in this cluster were early-responsive to cold stress, indicating that the ethylene-mediated stress response is activated as soon as cold stress is applied to rubber trees. Four of the early-responsive enzymes were annotated as RING-type E3 ubiquitin transferases: Arabidopsis Tóxicos en Levadura - ATL2 and RING1, with two sequences each. ATL proteins are involved in the regulation of several plant stress-responsive genes (Serrano et al. 2006; Bopopi et al. 2010; Song et al. 2016, 2022), and RING1 proteins are necessary for the activation of programmed cell death (PCD) during the plant hypersensitive response (Lin et al. 2008; Lee et al. 2011).

The exclusive connection of two clusters, c14 and c159, indicated a direct mechanistic association. Two early-response genes in this cluster were annotated: a hub gene that corresponded to a nematode resistance protein-like HSPRO2 and BON1-associated protein 2 (BAP2), a gene that was connected to two c14 genes. BAP proteins are negative regulators of cell death in plants and are essential for plant development and the hypersensitive response, and their expression is modulated by temperature (Yang et al. 2007; Hou et al. 2018; Cao et al. 2019). Other genes involved in PCD regulation were also present in other clusters. In fact, the PCD process appeared to be a current process throughout the rubber tree short-term cold response. Cluster c101 included a regulator of PCD: Myb108. Myb108 is a negative regulator of ABA biosynthesis and ABA-induced cell death (Cui et al. 2013) and is highly induced in rose plants under chilling (4°C) and freezing (-20°C) treatments. Arabidopsis plants overexpressing the rose ortholog showed improved cold tolerance and better performance under other stresses as well as a shorter growth cycle than the wild type (Dong et al. 2021). Cluster c161 included both proteins that positively (RING1) or negatively (Accelerated cell-death ACD11) regulate cell death. ACD11 is a sphingolipid transfer protein that controls cell death through the regulation of sphingolipid levels (Brodersen et al. 2002; Simanshu et al. 2014). ACD11 transiently accumulates under low concentrations of ABA and under salt and drought stress, and its overexpression confers abiotic stress tolerance (Li et al. 2020). Sphingolipids constitute a significant part of plant plasma membrane lipids and are also involved in mediating SA signaling in the plant stress response (Pata et al. 2010; Alden et al. 2011; Vicente et al. 2012), and cluster c126 appeared to manage sphingolipid metabolism. Finally, cluster c194 had proteins involved in the negative regulation of SA-mediated PCD (MACPF domain-containing proteins CAD1 and NSL1) (Noutoshi et al. 2006; Tsutsui et al. 2006). Considering the probable functions of these proteins and connections within the module, the results indicated that rubber trees trigger the PCD process early in cold exposure and maintain tight regulation of the mechanism over time.

Cluster c159, connected with c53 and c143, formed a network with clusters c34, c121 and c161 to promote the nucleotide sugar metabolic pathway. In addition, in module (iii), all three clusters have genes involved in nucleotide sugar metabolism. Nucleotide sugars are essential for cell wall biosynthesis and can function as signaling molecules, being sugar donors for the targeted glycosylation of different compounds (Figueroa et al. 2021). These results suggested that one of the most important response cascades to cold stress in rubber tree involves nucleotide sugar signaling and cell wall biosynthesis activation.

In module (ii), cluster c34 was the main player in Ca^2+^ signaling. The c34 cluster was enriched for transmembrane transport of Ca^2+^, widely known as a second messenger in plants (Yuan et al. 2018; Michailidis et al. 2020; Zhang et al. 2021). Low temperature stimulus has been well established as a main trigger of cell Ca^2+^ influx by the activation of Ca^2+^ channels, which leads to the Ca^2+^-signaling cascade of the plant cold stress response (Mori et al. 2018; Liu et al. 2021; Mao et al. 2021). Several metabolites act as low-temperature sensors, after which cells release calcium through the MAPK signaling pathway. The MAPK signaling pathway triggers signal transduction via TFs such as ERF, bHLH and MYB (Sun et al. 2021). Because of the overrepresentation of proteins with Ca^2+^ transporter activity in cluster c34, it could be considered the cluster controlling Ca^2+^-influx, therefore regulating Ca^2+^-signaling, which agrees with the cluster central position in the upregulated module (ii).

Cluster c143 was also connected to cluster c28. In addition to being enriched for stress responses and developmental processes, the GO terms “Positive regulation of protein kinase activity” and “DNA-binding transcription factor activity” were also overrepresented GO terms in the c28 cluster. A transcript (PASA_cluster_65283) stood out in cluster c28 for having the second highest BC and SC values in the whole GCN; however, there was no functional annotation for the sequence, making it a strong candidate for further analysis into the rubber tree cold stress response. Five of its connected transcripts were annotated, and the proteins are involved in plant meiotic recombination (Homologous-pairing protein 2 homolog, c143) (Uanschou et al. 2013; Shi et al. 2019), zeaxanthin synthesis (Beta-carotene 3-hydroxylase 1, chloroplastic, c106) (Fiore et al. 2006), JA-dependent defense response (Suppressor of NPR1-1 Constitutive 4, c28) (Bi et al. 2010), stress-induced osmotic solute accumulation (Probable polyol transporter 4, c28) (Noiraud et al. 2001) and Ca^2+^-signaling (Probable calcium-binding protein CML36, c28) (Astegno et al. 2017).

Additionally, linked to cluster c28, presenting only this connection in the HCCA network, cluster c126 exhibited enrichment for the stress response. Phenylalanine, apart from its role in protein biosynthesis, is the precursor of several plant phenolic compounds, such as lignin, flavonoids and anthocyanins (Maeda and Dudareva 2012), and salicylic acid (SA) (Lefevere et al. 2020). The remaining five clusters in module (ii) did not show any overrepresentation of GO terms in the enrichment analysis; nevertheless, the genes included in these clusters are involved in signaling processes, response to stress and transcription regulation. Together with clusters c28 and c34, cluster c121 had genes involved in the ethylene biosynthesis pathway and was the cluster with the most enzymes mapped to the KEGG pathways. Cluster c101 had the action of BAM1, a receptor protein kinase involved in plant development (DeYoung et al. 2005) that also has a role in the drought stress response and is required for long-distance signaling for stomatal closure (Takahashi et al. 2018). Cluster c194 included sugar transporters involved in cell wall biogenesis (UDP-galactose transporter 2) (Norambuena et al. 2005) and plant growth and stress response (Sugar transport protein 13) (Schofield et al. 2009; Lee and Seo 2021), in addition to having lignin synthesizing proteins.

### 3.2 Identification of Key Elements within Cold Response-Associated Modules

In addition to evaluating the functional profile of all the associated genes within the modules selected, we performed inferences into the importance of the genes with high centrality within a GCN - hubs, which may be described as core regulators (Amrine et al. 2015). Through the criteria defined for selecting hubs, we identified each cluster of the cold stress gene modules, which were considered key elements in the cold resistance definition (Carlson et al. 2006; Koido et al. 2018).

Among the nineteen hub genes of the downregulated module (i), two had functional annotation indicating a downregulation of photosynthetic pathway genes (clusters c42 and c166). The c42 hub gene was annotated as lycopene beta cyclase, chloroplastic/chromoplastic (LCYB), an enzyme of the carotenoid biosynthesis process. This hub highlighted the importance of the formation of pigment-protein complexes in the photosystems protecting the machinery from reactive oxygen species (ROS) created by abiotic stresses (Dall’Osto et al. 2006; Shi et al. 2015; Kang et al. 2018). The hub from cluster c166 was annotated as OTP51, a pentatricopeptide repeat-containing protein that is required for the proper assembly of photosystems I and II in Arabidopsis and rice (de Longevialle et al. 2008; Ye et al. 2012). Combined with low temperature, light promotes an enhancement in the transcription of cold-responsive and photosynthesis-related genes (Soitamo et al. 2008). The downregulation of both genes in rubber trees might be linked to the reduction in photosynthetic activity due to cold stress; nevertheless, this modulation was detected in the dark period of the experiment.

In addition to the regulation of photosynthetic activity from the cold-downregulated genes, it was also possible to identify two hubs representing proteins involved in abscisic acid signaling, which is essential for development and responses to abiotic stress. A transcript similar to protein phosphatase 2C (PP2C) was one of the hubs in cluster c171. Chao et al. (2020), in a genome-wide identification and expression analysis of the phosphatase 2A family in rubber trees, identified cis-acting elements related to the low temperature responsiveness (LTR) category. Under non-stress conditions, PP2C proteins negatively regulate the ABA-mediated signaling pathway by inactivating ABA-responsive genes. Once under stress, ABA receptors bind PP2Cs and inhibit their activity, consequently activating ABA signals (Cai et al. 2017). The other hub gene belonged to cluster c134 and was annotated as zinc finger CONSTANS-like 4 protein (COL4), a transcription factor. COL4 is a flowering inhibitor that represses flowering locus T (FT) gene expression (Steinbach 2019), and strikingly, it is also involved in abiotic stress responses through the ABA-dependent signaling pathway in Arabidopsis (Min et al. 2014). Considering that ABA concentration might have increased in the rubber trees during the cold exposure, which is enforced by the downregulation of a transcript similar to a PP2C protein, one could expect this gene to be upregulated instead. ABA is known to delay flowering; nevertheless, under severe drought conditions, ABA upregulates florigen genes, which is part of the drought escape (DE) strategy. DE prompts plants to accelerate their vegetative growth and reproduction stages during the period of high water availability. In the drought season, the plants enter a dormant stage (Verslues and Juenger 2011; Yıldırım and Kaya 2017).

The downregulation of three other strongly coexpressed hub genes, directly connecting three different clusters (c184, c171 and c153), identified in the last phase of the cold experiment also suggested an increase in ABA synthesis by the *H. brasiliensis* plantlets connected to an inhibition of growth and the photoperiod. The hub identified for cluster c184 was annotated as homeobox-leucine zipper (HD-Zip) protein HAT5 (AtHB1), which was connected to the PP2C hub, and it was identified as a downregulated gene after 12 hours of rubber tree cold exposure. HD-Zip transcription factors appear to be a plant-specific TF family, and their members take part in plant development and stress responses (Ariel et al. 2007). Each member has its own expression pattern under different environmental conditions (Li et al. 2019, 2020). This hub was connected not only to the c171 hub but also to the c153 hub, which was annotated as a leaf rust receptor-like kinase 10-like (LRK10L) protein that has an Arabidopsis homolog gene involved in ABA signaling (Lim et al. 2014).

The hub from cluster c156 was annotated as histone acetyltransferase of the MYST family (HAM) 1/2 proteins, which are involved in UV-B DNA damage repair (Campi et al. 2012). Interestingly, these proteins positively regulate the expression of the FLOWERING LOCUS C (FLC) gene, therefore being a flowering repressor as well (Xiao et al. 2013). The downregulation of this transcript in *H. brasiliensis* plantlets might be due to a decrease in DNA damaged by UV-B because of the absence of light. In addition, the downregulation of the c196 hub annotated with the Down Homeobox protein BEL1 homolog from different metabolic processes that are flowering repressors (Bhatt et al. 2004) suggested that rubber trees may have a DE-type strategy to deal with abiotic stresses.

Eighteen hub genes were identified in the upregulated module (ii). Five hub genes were annotated as Ferrochelatase-2 (FC2, c121), SNAP25 homologous protein SNAP33 (SNAP25, c34), Ubiquinol/Alternative oxidase (AOX, c121), Endochitinase (CHI, c121) and Heat shock cognate 70 kDa protein 2 (HSP70-2, c101) were strongly coexpressed in the light period of the experiment. FC2 is involved in photosynthetic cytochrome biogenesis and positively regulates chlorophyll and carotenoid contents (Scharfenberg et al. 2014), while AOX maintains the energy balance between mitochondria and chloroplasts by maintaining coordination between the activities of both organelles and is generally upregulated in response to cold stress (Vanlerberghe 2013; Vanlerberghe et al. 2016). Photosynthesis and respiration are coupled into carbon and energy metabolism, and both processes generate ROS. Although ROS accumulation has a deleterious effect on cells, ROS are also stress-signaling molecules. As such, in addition to protecting the photosynthesis and respiration complexes, there is a modulation of the stress-signaling pathways (Vanlerberghe 2013). The expression of the SNAP25 protein is upregulated by pathogen infection and mechanical stress (Wick et al. 2003), and the protein is also involved in the drought stress response (Nisa et al. 2017; Wang et al. 2017). SNAP25 is a protein unit of the SNARE protein complex, which is essential for vesicle trafficking. A CHI protein is tightly coexpressed with SNAP25. Endochitinases are known as pathogen-related defense responses, but their induction is generally nonspecific, being enhanced by both biotic and abiotic stresses and hormones, such as ethylene. They can either function inside the cell or be secreted (Collinge et al. 1993; Cletus et al. 2013). Together with the protection of energy levels and augmented stress-related vesicle trafficking, a chaperone of the HSP70 family protein exerts a high influence in module (ii). HSP70 proteins depend on ATP to properly function, and they are required for the transport of proteins to chloroplasts and mitochondria (Mirus and Schleiff 2009). In Arabidopsis, *HSP70-2* is involved in *FLC* expression regulation, hormone signaling pathways and abiotic stress responses, including responses to cold. Members of its family have redundant functions, but double and triple mutants presented accelerated growth, branching and flowering (Leng et al. 2016).

HSPRO2 protein is a positive regulator of basal disease resistance and regulates plant growth through interaction with the SnRK1 complex. The protein might also be involved in the leaf senescence process (Gissot et al. 2006; Schuck et al. 2013). *HSPRO2* expression was upregulated in rubber tree in the early stage of cold exposure and was highly correlated with three other hub genes in cluster c159; nevertheless, these genes did not have a functional annotation. Understanding the importance of these unknown protein genes in a cluster enriched for stress response processes in rubber tree requires exploring how these genes might affect cold stress adaptation in *Hevea*. These genes can also be candidates for analysis in other plant species since they are present in their genomes.

One of the early-responsive *ATL2* genes in cluster c14 was also a hub gene. In Arabidopsis, *ATL2* is upregulated after treatment with defense response elicitors (Serrano et al. 2006). The overexpression of the poplar homolog in tobacco promoted the upregulation of defense - and PCD-related genes, indicating a role in ubiquitination-mediated regulation of the defense response (Bopopi et al. 2010). Interestingly, the potato *ATL2* homolog negatively regulates *COR* gene expression, possibly by content-dependent ubiquitination of CBF proteins (Song et al. 2022). Another early-responsive gene was a hub in cluster c14, and it was annotated as a probable galacturonosyltransferase-like 10 (GATL10) protein. GATLs are required for plant primary cell wall biosynthesis (Kong et al. 2009, 2011, 2013). Rice and eucalyptus *GATLs* are upregulated by several types of abiotic stresses (Liu et al. 2016; Cheng et al. 2018). Plant cell wall composition and cell wall-modifying enzyme activities change in response to cold stress. Enhanced lignin synthesis is one of the strategies to cope with chilling stress by strengthening the cell wall and preventing cellular damage (Le Gall et al. 2015).

Considering all these key elements of transcriptional regulation of rubber trees in response to cold, we were able to characterize the general genetic responses to cold stress of two important clones of H. *brasiliensis*, which serve as a basis for rubber breeding, using GCN. As a tropical tree species, rubber tree most likely did not encounter prolonged periods of low temperatures during its evolution; nevertheless, the few wild genotypes that were successfully introduced in Southeast Asia carried such much genetic diversity that breeding for varieties that are cold tolerant is still possible. Using the GCN strategy applied in this study, we were able to visualize *Hevea*’s primary reprogramming of gene expression and the relationship among the genes involved in the cold stress response. In the short period of cold exposure studied in this work, the plantlets activated the ethylene-mediated signaling pathway from the beginning of the stress treatment and kept ethylene signaling active. Programmed cell death played a major role in the rubber tree cold response process and is tightly regulated by signaling cascades. Growth inhibition and cell wall thickening are implemented by the plantlets, which can be correlated to a possible drought escape strategy triggered by cold stress. In view of the genotypes analyzed, our results may represent the species’ genetic stress responses developed during its evolution. The understanding of how *H. brasiliensis* copes with low-temperature stress can greatly improve the breeding strategies for this crop and emphasize the importance of rubber tree genetic diversity preservation since it has such a narrow genetic base, is being impacted by climate change and is the only source for large-scale rubber production.

## 4 Methods

### 4.1 RNA-Seq Experiment

In this study, two genotypes of *H. brasiliensis* recommended for escape areas (Mantello et al. 2019), GT1 and RRIM600, were evaluated. Both genotypes were collected at the Agência Paulista de Tecnologia dos Agronegócios/SAA, Votuporanga, São Paulo, Brazil (de Souza et al. 2015). In low temperatures, RRIM 600 presents an avoidance physiological strategy, while GT1 balances photosynthetic/growth activities and the stress response (Mai et al. 2010; Mantello et al. 2019). The cold experiment consisted of exposing three six-month-old seedlings of each genotype to a temperature of 10°C for 24 hours (12 h light/12 h dark). During this period, leaf material was collected in a time series at 0 hours (0 h - control), 90 minutes (90 m), 12 hours (12 h), and 24 hours (24 h). The RNA extracted from these samples was used to construct cDNA libraries, and it was subsequently sequenced using an Illumina Genome Analyzer IIx with the TruSeq SBS 36-Cycle Kit (Illumina, San Diego, CA, USA) for 72 bp PE reads. The transcriptome assembled by Mantello et al. (2019) was used for the analyses in the present study.

### 4.2 Bioinformatics and Differential Gene Expression Analysis

The reads of each genotype evaluated with their respective times and replicates of treatment were mapped with the Bowtie2 aligner (Langmead and Salzberg 2012) in the reference genome (Tang et al. 2016). The estimated abundance of reads for each sample was calculated by RSEM (Li and Dewey 2011), and subsequently, a matrix was constructed with expression values and abundance estimates between samples and replicates. Genes with a minimum of 10 counts per million (CPM) in at least three samples were retained for further analyses. These samples were normalized by the quantile method and transformed into log2 counts per million (log2cpm) using the edgeR package (Robinson et al. 2010).

From the normalized data, a principal component analysis (PCA) was performed using R software to explore the data and verify the behavior of the samples. For DEG analysis, the datasets from GT1 and RRIM600 were combined to create a unique *H. brasiliensis* expression dataset. An empirical Bayes smoothing method implemented in the edgeR package (Robinson et al. 2010) was used to identify DEGs between cold exposure sampling times, in which the following comparisons were performed: (i) 0 h vs. 90 m; (ii) 0 h vs. 12 h; (iii) 0 h vs. 24 h; (iv) 90 m vs. 12 h; (v) 90 m vs. 24 h; and (vi) 12 h vs. 24 h. The p-values were adjusted using a Bonferroni correction and genes with a p-value ≤ 0.05 were considered DEGs.

Rubber tree DEGs were compared between the time points evaluated with Venn plots constructed with the Venn R package (Dusa 2017). For all genes, we selected the respective annotation obtained with the Trinotate v2.0.1 pipeline ((Bryant et al. 2017); https://trinotate.github.io/). Untranslated transcripts were searched against the SwissProt/UniProt database using BLASTX, filtered with an e-value of 1e-5, and placed into a tab-delimited file. Transcripts were also associated with Gene Ontology (GO) (Harris et al. 2004).

### 4.3 Gene Coexpression Network (GCN) Analysis

The GCN was inferred through R Pearson correlation coefficients with a cutoff of 0.8 (van Noort et al.

2004). The R values of the GCN were normalized using the highest reciprocal rank (HRR) approach limited to the top 10 connections, and the heuristic cluster chiseling algorithm (HCCA) was used to organize the network into communities (Mutwil et al. 2010). Only the top 10 Pearson R correlation coefficients considering a 0.8 cutoff were retained for each gene, and the genes were not considered disconnected. The HRR networks and the HCCA results were visualized with Cytoscape software v.3.8.0 (Shannon et al. 2003).

### 4.4 Cold Stress Association

The number of DEGs in the clusters was used to pinpoint the most likely groups involved in the *Hevea* cold stress response. Hub genes in the selected clusters were analyzed using the node degree distribution (DD), betweenness centrality (BC), and stress centrality distribution (SC) parameters of the networks that were obtained using Cytoscape software v.3.8.0 (Shannon et al. 2003). Genes with high DD that presented high values of BC and SC were considered highly influential in the selected clusters’ coexpression dynamics. If a highly connected node had low BC and SC values, it was not selected as a hub. For each network cluster, at least one hub gene was identified, and the hub genes were evaluated by their functional annotation and/or the genes with which they interacted.

To identify molecular associations with the cold stress response, we selected the clusters that together accounted for 70.2% of the DEGs identified. We considered as associated clusters those that contained at least 50% of their genes as DEGs, expanding the module across these clusters’ neighbors considering a minimum required frequency of 20% of DEGs within a cluster (Table S6).

### 4.5 Functional Enrichment

Since functional coherence between coexpressed genes belonging to the same cluster is expected, the analysis of enrichment of GO terms was performed in each cluster identified by the HCCA. Overrepresented GO terms in the clusters were determined with BiNGO plugin v.3.0.4 (Maere et al. 2005) using a customized reference set created from the transcriptome annotation data. A hypergeometric distribution and false discovery rate (FDR) < 0.05 were used in the analyses. GO enrichment analyses were performed using EnrichmentMap plugin v.3.3.1 (Merico et al. 2010) with FDR Q-values < 0.05 and BiNGO output files. Both plugins were used in Cytoscape software v.3.8.0 (Shannon et al. 2003).

## Acknowledgments

We would like to acknowledge the Fundação de Amparo à Pesquisa do Estado de São Paulo (FAPESP), the Conselho Nacional de Desenvolvimento Científico e Tecnológico (CNPq), the Coordenação de Aperfeiçoamento de Pessoal de Nível Superior (CAPES) and the Instituto Agronômico de Campinas (IAC), for support in the field and technical and scientific support from Dr. Paulo de Souza Gonçalves.

## Statements and Declarations

### Ethics Approval

We confirm that no specific permits were required to collect the leaves used in this study.

### Consent to Participate

This work was a collaborative study developed by researchers from EMBRAPA (Brazil), IAC (Brazil), and UNICAMP (Brazil). We confirm that the field studies did not involve endangered or protected species.

### Consent for Publication

Not applicable

### Data Availability

The datasets analyzed for this study can be found in the NCBI database

(https://www.ncbi.nlm.nih.gov/bioproject/PRJNA483203) and are included in the Supplementary Information files.

### Authors’ Contributions

AS, CS, CM and RV conceived the project; CS, SB, AA, FF, RB and RV performed the analyses and wrote the manuscript. All authors reviewed, read, and approved the manuscript.

