## Additional Files Description for "A deeper cascade of mechanisms involved in cold resistance in *Hevea brasiliensis*" (from another version of this preprint: version 5, 2023-09-06)

**Supplementary information**

**Additional file 1: Table S1.** Gene Annotation. Annotation results for the 51,967 genes in the transcriptome used in this work include Gene Ontology terms, KEGG pathways, and protein names. (XLSX 1.87 Mb).

**Additional file 2: Table S2.** Differentially expressed genes across time periods. Comparison periods: 0 h vs. 90 min, 0 h vs. 12 h, 0 h vs. 24 h, 90 min vs. 12 h, 90 min vs. 24 h, 12 h vs. 24 h, 0 h+90 min vs. 12 h+24 h. (XLSX 75.2 kb).

**Additional file 3: Table S3.** Topological evaluation of genes within the modeled gene coexpression network. This table contains degree, betweenness, and stress values for all genes present in the GCN. (XLSX 1.16 Mb).

**Additional file 4: Table S4.** Topological evaluation of gene clusters within the major components of the gene coexpression network. This table contains degree, betweenness, and stress values for all clusters present in the major components of the GCN. (XLSX 14.3 kb).

**Additional file 5: Table S5.** GCN clusters, including differentially expressed genes. This table contains all clusters that include DEGs with information about the total number of genes, number of DEGs, proportion of DEGs, and selected groups. (XLSX 14.6 Kb).

**Additional file 6: Table S6.** Hub genes are identified in clusters within the established modules associated with the cold response. This table contains information about the selected modules, hub annotations, DEG conditions, degree, betweenness, and tree values for each hub gene. (XLSX 12.6 kb).
